# Force Field Limitations of All-Atom Continuous Constant pH Molecular Dynamics

**DOI:** 10.1101/2024.09.03.611076

**Authors:** Craig A. Peeples, Ruibin Liu, Jana Shen

## Abstract

All-atom constant pH molecular dynamics simulations offer a powerful tool for understanding pH-mediated and proton-coupled biological processes. As the protonation equilibria of protein sidechains are shifted by electrostatic interactions and desolvation energies, p*K*_a_ values calculated from the constant pH simulations may be sensitive to the underlying protein force field and water model. Here we investigated the force field dependence of the all-atom particle mesh Ewald (PME) continuous constant pH (PME-CpHMD) simulations of a mini-protein BBL. The replica-exchange titration simulations based on the Amber ff19sb and ff14sb force fields with the respective water models showed significantly overestimated p*K*_a_ downshifts for a buried histidine (His166) and for two glutamic acids (Glu141 and Glu161) that are involved in salt-bridge interactions. These errors (due to undersolvation of neutral histidines and over-stabilization of salt bridges) are consistent with the previously reported p*K*_a_’s based on the CHARMM c22/CMAP force field, albeit in larger magnitudes. The p*K*_a_ calculations also demonstrated that ff19sb with OPC water is significantly more accurate than ff14sb with TIP3P water, and the salt-bridge related p*K*_a_ downshifts can be partially alleviated by the atom-pair specific Lennard-Jones corrections (NBFIX). Together, these data suggest that the accuracies of the protonation equilibria of proteins from constant pH simulations can significantly benefit from improvements of force fields.

## Introduction

Solution pH mediates many important biological processes through coupling proton titration with conformational changes of proteins. Over the last two decades, constant pH methods have been developed to rigorously account for solution pH in condensed-phase molecular dynamics (MD) simulations. Instead of fixing the protonation states of protein sidechains to the initial conditions, e.g., histidines with one proton on either the *δ* or *ϵ* imidazole nitrogen, constant pH simulations allow protonation states to respond to changes in the electrostatic environment accompanying conformational dynamics. Currently, the main approaches to enable proton-coupled dynamics is through Monte-Carlo (MC) sampling of protonated and deprotonated states (discrete or hybrid MD/MC constant pH methods)^1–5^ and an extended Hamiltonian description whereby an auxiliary set of fictitious particles are propagated to represent proton titration (continuous constant pH or *λ* dynamics based constant pH methods).^6–11^ A more detailed discussion of the development of constant pH methods is given in a recent review^12^ for more references.

In the early developments of constant pH methods, various generalized Born (GB)^2,6,7,13^ or Poisson-Boltzmann (PB) implicit solvent models are used for both conformational and protonation state sampling. The so-called hybrid-solvent scheme combines the conformational sampling in explicit solvent with implicit-solvent calculation of titration energies^4,14^ or forces,^8^ resulting in significantly improved accuracies of the calculated p*K*_a_ values. The more recent developments^11,15–18^ utilize explicit solvent for both conformational and protonation state sampling, allowing constant pH simulations of systems that implicit-solvent representations are less accurate for, e.g., nucleic acids,^19,20^ surfactant micelles,^21^ and proteins in lipid bilayers,^22^ although careful reparameterization of dielectric boundary has mitigated some of the issues. Perhaps the most significant drawbacks of using continuum models (either fully implicit-solvent simulations or with the hybrid-solvent scheme) are the failure to account for explicit interactions with ions^23^ and the poor computational scaling behavior, *n*^2^, where *n* is the number of solute atoms, although work of Onufriev and coworker demonstrated that *nlogn* can be achieved through a Hierarchical Charge Partitioning (HCP) approximation.^24^ The first all-atom *λ* dynamics based CPHMD^MS*λ*D^ implementation in CHARMM was applied to calculate the p*K*_a_’s of RNAs^25^ and peptides inserted in the lipid bilayer.^26^

By removing the aforementioned deficiencies due to the implicit-solvent models, all-atom constant pH simulations can in principle (i.e., given converged simulations) offer more accurate p*K*_a_ values than implicit- or hybrid-solvent constant pH simulations. The benchmark simulations using the first all-atom particle mesh Ewald (PME) continuous constant pH MD (CpHMD) implementation in CHARMM^16^ demonstrated improved correlation between the calculated and experimental p*K*_a_ shifts relative to the solution (also known as the model) values as compared to its “predecessor”, the hybrid-solvent CpHMD in CHARMM,^8^ even though the root-mean-square errors (rmse) with respect to the experimental data are similar.

The key physical determinants of protein p*K*_a_ shifts relative to the model values are electrostatic interactions and desolvation free energies, which have been shown to vary between different protein force fields for proteins and water.^27–32^ As such, p*K*_a_ values derived from constant pH simulations can be used to probe the force field deficiencies.^33^ So far, the benchmark simulations of all-atom constant pH methods^15–18,34^ have been conducted using the CHARMM c22/CMAP,^35,36^ c36,^37^ or c36m^38^ force fields, while other force fields such as the widely used Amber ff14sb^39^ or ff19sb^40^ force fields have not been tested. Recently, Sequeira et al. compared the p*K*_a_ calculations using the hybrid-solvent MD/MC constant pH simulations with the GROMOS 54A7 and CHARMM c36m force fields.^41^ Their study^41^ demonstrated significant difference between the two force fields in calculating the highly shifted p*K*_a_’s of Asp26 in human thioredoxin and Cys32/Cys35 in *E. coli* thioredoxin. Specifically, the c36m calculated p*K*_a_’s were very stable over the simulation time (up to 50 ns), whereas the 54A7 calculated p*K*_a_’s converged much slower but ultimately achieved better agreement with experiment.

In this work, we investigate the force field dependence of the all-atom PME CpHMD simulations by comparing the titration simulations of a mini-protein BBL (Figure 1) based on the Amber ff19sb,^40^ ff14sb,^39^ and CHARMM c22/CMAP^35,36^ protein force fields and their respective preferred water models, TIP3P,^42^ OPC,^43^ and CHARMM-style TIP3P.^35^ The p*K*_a_ data based on the c22 force field are taken from our previous work.^34^ BBL is a well suited benchmark protein for the constant pH based p*K*_a_ calculations, as it contains buried histidines and carboxylic acids involved in salt-bridge interactions, for which large p*K*_a_ shifts relative to solution (also called model) values are expected and challenging to predict accurately.^44^ Also, BBL has been previously used to benchmark the performance of the all-atom PME CpHMD implementations^16,34^ in CHARMM^45^ and Amber programs.^46^ Importantly, due to its small size, conformational sampling is less likely an issue. We found that the largest error for all force fields is for a buried histidine, while glutamic acids involved in salt bridge interactions have overestimated p*K*_a_ shifts. Specific ion binding was observed in the ff19sb simulations; however, the use of atom-pair specific corrections to the Lennard Jones parameters (widely known as NBFIX)^47–51^ did not reduce the errors of the calculated p*K*_a_’s.

**Figure 1:**
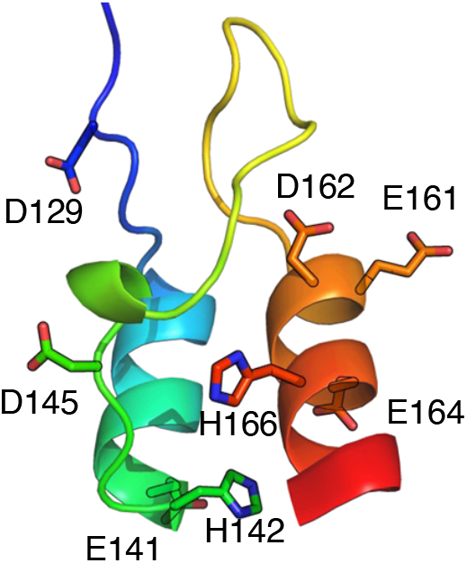
Structure and titratable sites of BBL. The NMR model of BBL (PDB ID: 1W4H,^52^ first entry) The titratable sidechains are labeled and shown in the stick model.

## Methods and Protocols

Unless otherwise noted, the simulations were performed using the all-atom particle-mesh (PME) CpHMD implementation^34^ in Amber24.^46^

### Preparation of the protein system

The coordinates of BBL were retrieved from the Protein Data Bank (PDB) entry 1W4H.^52^ The first NMR model was used. The NMR model shows no interactions between the terminal groups and any titratable residues. For simulations, the N- and C-termini were acetylated (ACE-) and amidated (-NH2), respectively. Following our previous protocol,^53^ the positions of hydrogens were built using the HBUILD command in the CHARMM program,^45^ and a custom CHARMM script was used to add dummy hydrogens to the *syn* positions of carboxylate oxygens on all Asp and Glu sidechains. Next, the CHARMM coordinate file was converted to the Amber format. The protein was solvated in a truncated octaheral water box with a minimum of 15 Å between the protein heavy atoms and the water oxygens at the box edges. To neutralize the simulation box at pH 7.5 and reach an ionic strength of 150 mM, 18 Na^+^ and 19 Cl^−^ were added. Note, to accommodate the replica-exchange protocol, the number of counter ions was not adjusted for other pH conditions (see Concluding Discussion).

### Minimization, heating, and equilibration of the protein

The *pmemd*.*cuda* engine of the Amber2024 program^46^ was used for simulations. First, the protein system underwent 10,000 steps of energy minimization using the steepest descent (1000 steps) and conjugate gradient (9000 steps) algorithms, whereby the protein heavy atoms were harmonically restrained with a force constant of 100 kcal·mol^*−*1^ Å ^*−*2^. Next, the PME-CpHMD module^34^ was turned on and the restrained system was heated from 100 to 300 K over 1 ns at pH 7.5 with a 1 fs timestep in the NVT ensemble. Finally, a two-stage equilibration was performed in the NPT ensemble. In the first stage, the PME-CpHMD simulation was performed at pH 7.5 with a 1-fs time step. A harmonic restraint was placed on heavy atoms, and it was reduced from 100 in the first 1 ns to 10 kcal·mol^*−*1^ Å ^*−*2^ in the second 1 ns. In the second stage, 16 pH replicas were placed with increments of 0.5 units in the pH range 1.0–8.5 and simulated independently for 2 ns, employing a 2 fs timestep and harmonic restraints on the backbone heavy atoms. Here the restraints were gradually reduced in four 500-ps steps, from 10 to 5, 2.5, and 0 kcal*·*mol^*−*1^*·*Å ^*−*2^.

### Production pH replica-exchange CpHMD simulations

In the production run, the asynchronous pH replica-exchange (REX) protocol^54^ was turned on using the same 16 pH replicas as in the equilibration stage. The exchange between adjacent pH was attempted every 1000 MD steps (2 ps) according to the Metropolis criterion.^8^ The REX simulations were run for 32 ns per replica, with the aggregate simulation time of 512 ns. The p*K*_a_’s for all residues were converged (Supplementary Figure S1–S4). The first 10 ns per replica was discarded in the conformational analysis and p*K*_a_ calculations.

### MD protocol

The SHAKE algorithm was employed to constrain the bonds involving hydrogens whenever a 2-fs step was used. The production runs were performed in the NPT ensemble, where the temperature was maintained at 300 K using the Langevin thermostat with a collision of 1.0 ps^*−*1^, and the pressure was maintained at 1 atm using the Berendsen barostat with a relaxation time of 1 ps. The PME method was used to calculate long-range electrostatics with a real-space cutoff of 12 Å and a 1-Å grid spacing in the re-ciprocal space calculation. Consistent with the our previous titration simulations with the c22 force field,^34^ Lennard-Jones energies and forces were smoothly switched off over the range of 10 to 12 Å.

### Model peptide p*K*_a_ values and protein p*K*_a_ corrections

The model titration parameters for ff19sb and ff14sb simulations of Asp, Glu, and His were taken from the previous work.^34^ The validation simulations^34^ were conducted using titration simulations of penta-peptides ACE-AlaAlaXAlaAla-NH_2_ (X = Asp, Glu, or His) at independent pH conditions with an interval of 0.5 pH in the pH range 2–5.5 for Asp, 2.5–6 for Glu, and 4–8 for His. The simulations lasted 20 ns at each pH. The calculated p*K*_a_’s along with the NMR determined values are given in Table 1. Note, ff14sb and ff14sb^fix^ gave nearly identical p*K*_a_’s for Asp and Glu penta-peptides. Given the small deviations from the experimental p*K*_a_’s, the following post-corrections were made to the calculated protein p*K*_a_’s. ff19sb: Asp (+0.35), Glu (+0.33), His (+0.06). ff14sb and ff14sb^fix^: Asp (+0.16), Glu (+0.18), His (−0.23). This is a caveat, as the corrections are different for individual residue types; the more rigorous way is to offset the model p*K*_a_’s before simulations. However, given the small magnitude of corrections (below 0.35 units), the resulting inaccuracies are negligible. Furthermore, for convenience, we retained the original pH scales in plotting the residue-based pH-dependent analysis.

**Table 1:** Calculated model peptide p*K*_a_’s based on Amber ff19sb and ff14sb force fields^*a*^.

|  | Asp | Glu | His |
| --- | --- | --- | --- |
| ff19sb | $3.32 \pm 0.07$ | $3.92 \pm 0.05$ | $6.48 \pm 0.06$ |
| ff14sb | $3.51 \pm 0.11$ | $4.07 \pm 0.07$ | $6.71 \pm 0.10$ |
| NMR | 3.67 | 4.25 | 6.54 |

<sup>a</sup>Titration simulations of penta-peptides ACE-AlaAlaXAlaAla-NH<sub>2</sub> (X = Asp, Glu, or His) were performed using the titration parameters from our previous work.<sup>34</sup> NMR derived $pK_a$ ’s are taken from Ref.<sup>55</sup>

We previously showed that protein p*K*_a_’s derived from the PME-CpHMD simulations are affected by the simulation box size, and the finite-size effect decreases with increasing number of water, becoming negligible with large water boxes.^34^ Since the current simulations employed a very large water box (15 Å cushion between protein and box edges), the calculated finite-size^16,34^ corrections were small (no larger than 0.1 unit). As such, no finite-size corrections were made to the calculated p*K*_a_’s.

### Force field parameters and CpHMD specific changes

The peptides and proteins were represented by the Amber ff19sb^40^ or ff14sb^39^ protein force field, with the respective OPC^43^ or TIP3P^56^ model for representing water. The ion parameters were taken Ref.^57^ For the simulations with ff14sb, the NBFIX corrections developed by Yoo and Aksimentiev (CUFIX) were applied to the atom pair specific Lennard-Jones parameters to destabilize the attractive interactions between Na^+^ and carboxylate (Asp^−^ or Glu^−^),^49^ between amine (Lys^+^) or guanidinium (Arg^+^) and carboxylate (Asp^−^ or Glu^−^).^50,51^ The all-atom PME CpHMD implementations in Amber include two force field modifications common to constant pH methods.^2,13,34^ First, to allow the single reference concept in constant pH simulations, the partial charges on the backbone are fixed to the values of a single protonation state (Asp^−^ /Glu^−^ and Hid/Hie), and the residual charge (ranging from 0.10 to 0.14 e for Asp, Glu and His) is absorbed onto the C-*β* atom for titration dynamics.^13^ Second, the rotation barrier around the carboxylate C-O bond is increased to 6 kcal/mol to prevent the dummy proton from rotating from the syn (initial position in the set up) to the anti position, in which case the proton would lose the ability to titrate.^2,13^

## Results and Discussion

### Comparison of the calculated p*K*_a_’s based on the Amber and CHARMM force fields and comparison to experiment

BBL has 6 carboxylic acids and 2 histidines. In this work, the calculated p*K*_a_’s based on ff19sb^40^ gave a root-mean-square error (rmse) of 1.26 pH units with respect to experiment, which is much larger than the rmse (0.62) from our previous PME-CpHMD simulations^34^ based on CHARMM c22,^35,36^ as well as the rmse (0.66) from our previous^13^ GBNeck2^60^ simulations with ff14sb^39^ (Table 2). Convergence and titration curves of ff19sb simulations are given in Supplementary Figure S1–S4. Note, the rmse of the p*K*_a_’s calculated from the preliminary ff14sb (without NBfix) simulations is 1.79, similar to ff19sb (Supplementary Table S1). Curiously, for both ff19sb and c22 simulations, the largest p*K*_a_ calculation error is for His166, which has the respective calculated p*K*_a_’s of 2.4 and 4.2 (Table 2). Both ff19sb and c22 simulations correctly predicted the p*K*_a_ downshift; however, the magnitude is too large compared to experiment (Table 2). For the ff14sb force field, the downshift is also overestimated (Supplementary Table S1). Without His166, the rmse for the ff19sb and c22 simulations are reduced to 0.73 and 0.48, respectively (Table 2), while for ff14sb the rmse without His166 is 1.38 (Supplementary Table S1). By contrast, the GB-Neck2 implicit-solvent simulations underestimated the p*K*_a_ downshift of His166 by 0.6.

**Table 2:** Comparison of the calculated p*K*_a_’s of BBL using the all-atom PME CpHMD titration based on Amber and CHARMM force fields.

| Residue | Expt | ff19sb<sup>a</sup> | ff14sb<sup>fixa</sup> | c22<sup>b</sup> | ff14sb/GB<sup>c</sup> |
| --- | --- | --- | --- | --- | --- |
| D129 | 3.9 | 3.5 | 3.6 | 3.6 | 2.8 |
| **E141<sup>d</sup>** | 4.5 | 3.4 | 3.1 | 4.0 | 4.1 |
| H142 | 6.5 | 6.6 | 6.5 | 5.8 | 6.9 |
| D145 | 3.7 | 3.3 | 2.2 | 3.2 | 2.6 |
| **E161<sup>d</sup>** | 3.7 | 2.6 | 2.6 | 4.0 | 3.3 |
| **D162<sup>d</sup>** | 3.2 | 3.3 | 1.7 | 2.9 | 3.2 |
| E164 | 4.5 | 3.6 | 3.4 | 4.0 | 4.0 |
| **H166<sup>d</sup>** | 5.4 | 2.4 | 2.1 | 4.2 | 6.0 |
| maxe<sup>e</sup> |  | -3.0 | -3.3 | -1.2 | -1.1 |
| rmse<sup>e</sup> |  | 1.26 | 1.58 | 0.62 | 0.66 |
| rmse (w/o H166) |  | 0.73 | 1.14 | 0.48 | 0.67 |

<sup>a</sup> This work. ff19sb refers to the ff19sb protein force field<sup>40</sup> with the OPC water model<sup>43</sup> and Amber default ion force field.<sup>57</sup> ff14sb<sup>fix</sup> refers to the ff14sb protein force field<sup>39</sup> with TIP3P water model<sup>56</sup> and the NBFIX corrected ion force fields.<sup>49–51</sup> <sup>b</sup> Taken from the previous work<sup>34</sup> using CHARMM c22 protein force field<sup>35</sup> with the CHARMM style TIP3P water model<sup>35,56</sup> and the NBFIX corrected ion force fields.<sup>48,58,59</sup> <sup>c</sup> Taken from the previous work<sup>13</sup> using the ff14sb protein force field and GBNeck2 implicit-solvent model.<sup>60</sup> <sup>d</sup> Residues discussed in the main text. <sup>e</sup> Maximal error (maxe) and root-mean-square error (rmse) are calculated with respect to the experimental data.<sup>61,62</sup>

Following His166, the largest p*K*_a_ calculation errors with the ff19sb force field are for Glu141 and Glu161, which have the calculated p*K*_a_’s of 3.4 and 2.6, compared to 4.0 and 4.0, respectively, from the c22 simulations (Table 2). Compared to the experimental p*K*_a_’s of 4.5 and 3.7 for Glu141 and Glu161, the ff19sb simulations underestimated both p*K*_a_’s by about 1.1 unit, whereas the c22 simulations underestimated the p*K*_a_’s by 0.5 and 0.3 units, respectively. Below we discuss the origins of the underestimated p*K*_a_’s for Glu141, Glu161, and His166.

### p*K*_a_ downshifts of Glu141 and Glu161 are due to salt bridge formation

The p*K*_a_’s of Glu141 and Glu161 calculated by the ff19sb simulations are both downshifted with respect to the model value, suggesting that they are involved in attractive electrostatic interactions. Trajectory analysis shows that Glu141 occasionally forms a salt bridge with Arg137, with the occupancy (or probability) increasing from zero at pH 1 to a maximum of nearly 20% near pH 4 where deprotonation is completed (Figure 2a–c). Over the same pH range, the solvent exposure of Glu141, defined by the number of water molecules in the first solvent shell of the carboxylate oxygens, also increases and plateaus to about 7 upon complete deprotonation (Figure 2d). Since the first solvent shell of carboxylate oxygens of the model pentapeptide (AAEAA) contains an average of 7 water molecules (SI), these analyses (Figure 2c,d) suggests that Glu141 becomes fully exposed to solvent once the salt bridge is disrupted.

**Figure 2:**
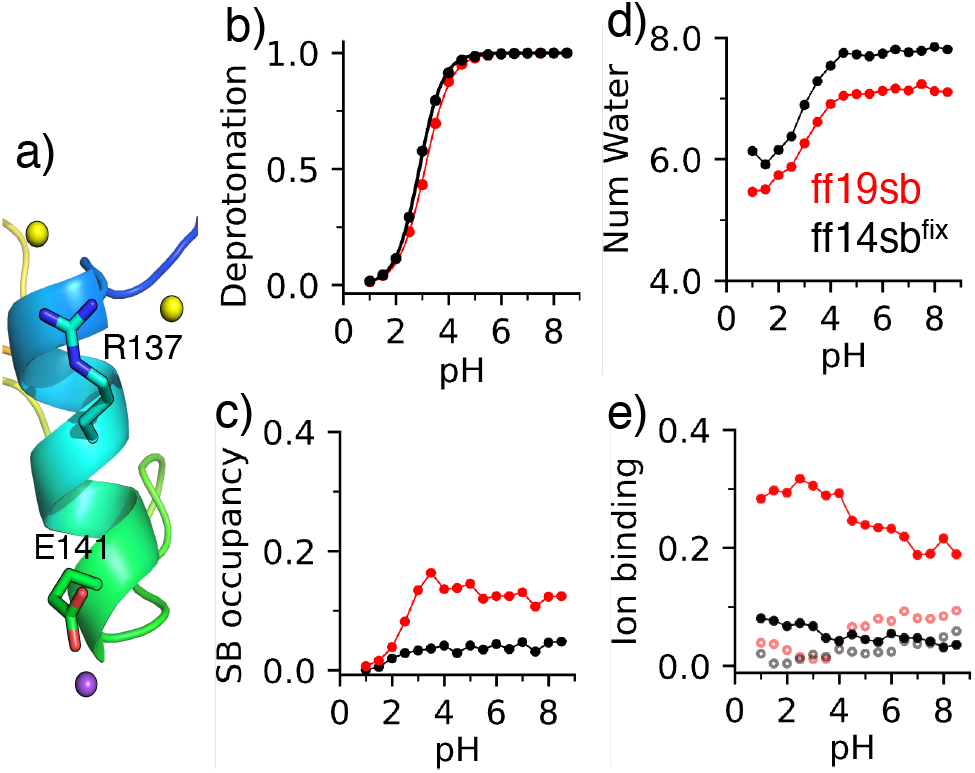
Titration of Glu141 in BBL and the pH-dependent salt bridge formation, solvent exposure, and ion binding. a) Snapshot of the bound ions from the ff19sb trajectory at pH 6.5. Na^+^ and Cl^−^ ions are shown as purple and yellow spheres, respectively. b) pH-dependent deprotonated fraction of Glu141. c) pH-dependent occupancy of the salt bridge formation between Glu141 and Arg137. A salt bridge is considered formed if the distance between a carboxylate oxygen of the charged Glu141 and a guanidinium nitrogen is below 4.0 Å. d) Number of water molecules within 3.4 Å of a carboxylate oxygen of Glu141 at different pH. e) Occupancy of Cl^−^ binding of Arg137 (closed circles) and Na^+^ binding of Glu141 (open circles) at different pH. An ion is considered bound if it is within a cutoff distance, 4.2 Å for Cl^−^ and 4.0 Å for Na^+^, from a guanidinium nitrogen of Arg137 or a carboxylate oxygen of Glu141. The data based on the ff19sb and ff14sbfix force fields are shown in red and black, respectively, and do not include p*K*_a_ corrections.

The strong correlation between the sigmoidal shaped pH profiles of the deprotonation fraction, salt-bridge occupancy, and solvent exposure of Glu141 indicates that the salt-bridge interaction, which stabilizes the charged state, is the main determinant for the p*K*_a_ downshift. This phenomenon has been observed in the previous titration simulations using the CpHMD implementations in the CHARMM and AMBER programs with the c22 force field.^16,34^ However, since the experimental p*K*_a_ of Glu is 4.5, which is 0.3 up shifted relative to the model value, the Glu141–Arg137 salt bridge interaction is likely overly stabilized by the ff19sb force field, and the overstabilization is to a lesser extent with the c22/CMAP force field as the p*K*_a_ underestimation is 0.5 unit smaller.

We next examined Glu161, which shows a p*K*_a_ downshift that is 1.2 units too large compared to experiment (Table 2). Similar to Glu141, deprotonation of Glu161 is also strongly correlated with the salt-bridge formation (with either Lys165 or Arg160, Figure 3a– d). The salt-bridge occupancy of Glu161 increases to a maximum of 28% near pH 5 (Figure 3d), which is higher than Glu141 (Figure 2c). Over the same pH range, the number of water molecules in the first solvent shell of Glu161 increases to about 6.5 (Figure 3e), which is slightly lower than that for Glu141 (Figure 2d). The increased salt-bridge interactions and slightly decreased solvent exposure favor the charged state, which may explain the lower p*K*_a_ of Glu161 relative to Glu141.

**Figure 3:**
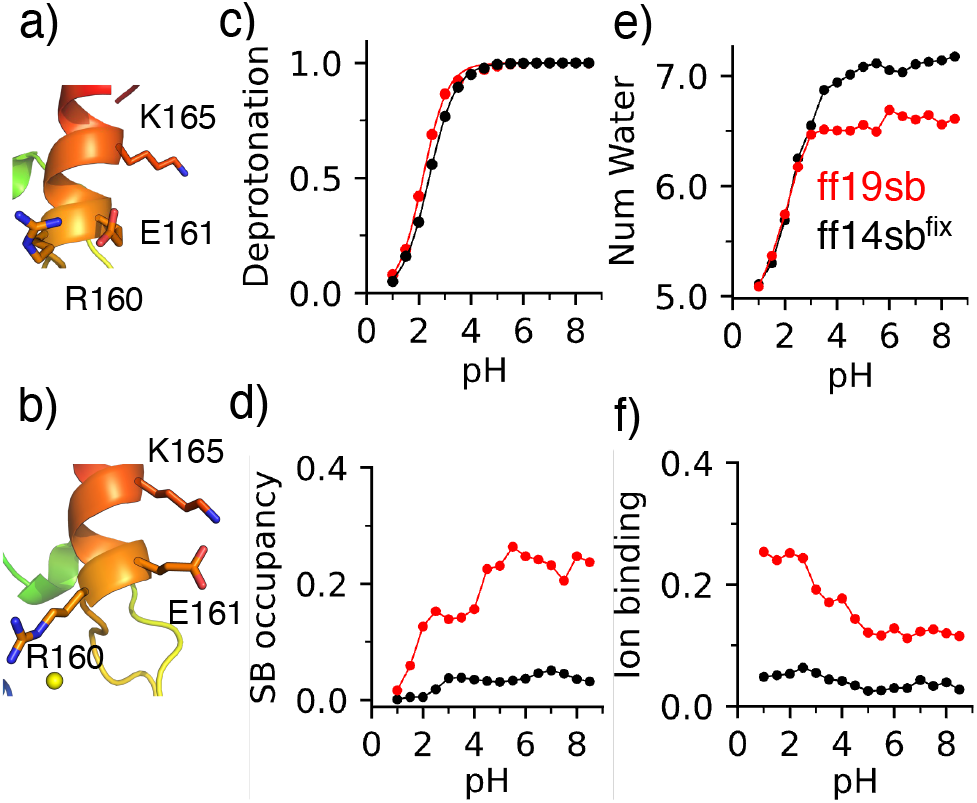
Titration of Glu161 in BBL and the pH-dependent salt bridge formation, solvent exposure, and ion binding. a) Snapshot showing a salt bridge between Glu161 and Arg160 taken from the ff19sb trajectory at pH 7. b) Snapshot of Arg160 bound to Cl^−^ and Glu161 forming a salt bridge with Lys165 from the ff19sb trajectory at pH 7. c) Deprotonated fraction of Glu161 at different pH. d) Salt bridge occupancy of Glu161 with either Arg160 or Lys165 as different pH. e) Number of water molecules within 3.4 Å of any carboxylate oxygen of Glu161 at different pH. f) Occupancy of Cl^−^ binding to Arg160 at different pH. The data based on ff19sb and ff14sbfix are shown in red and black, respectively, and do not include p*K*_a_ corrections.

### Downshifted p*K*_a_ of Asp162 is due to hydrogen bonding

The calculated p*K*_a_’s of Asp162 based on the ff19sb and c22 force fields are similar and downshifted relative to the model value by 0.4 and 0.7, respectively. These p*K*_a_ downshifts are similar to the experimental downshift of 0.5 units. Trajectory analysis suggests that the p*K*_a_ downshift of Asp162 can be attributed to the formation of a hydrogen bond (h-bond) with Thr152 (Figure 4a). The deprotonation of Asp162 occurs over the pH range of 1 to 5 (Figure 4b), where the increased deprotonation is accompanied by the h-bond formation between the carboxylate of Asp162^*−*^ and the hydroxyl group of Thr152 which increases from 5.3% to 83.8% (Figure 4c). Note, although protonated carboxylic can also serve as a h-bond acceptor to Thr152, the occupancy is very low (maximum *∼*20% at pH 2.5, Supplemental Figure S5). This phenomenon has also been observed in the c22 based PME-CpHMD simulations^16,34^ as well as the hybrid-solvent^63^ and implicit-solvent CpHMD simulations of a variety of proteins.^64^ The similarity between the experimental and calculated p*K*_a_’s of Asp162 based on the ff19sb and c22 force fields suggests that the h-bond interaction involving Asp162 may be represented appropriately by the force fields.

**Figure 4:**
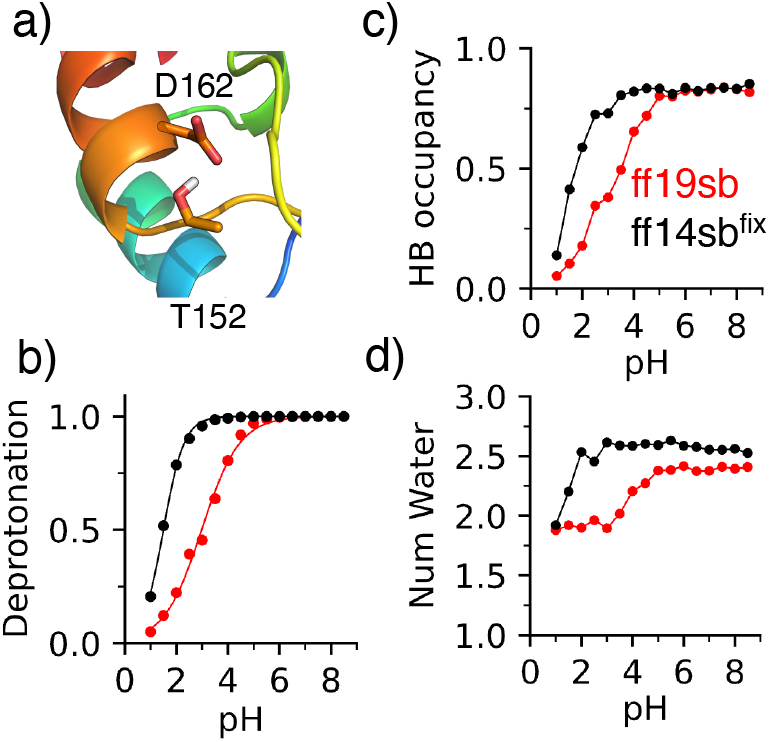
Titration of Asp162 in BBL and the pH-dependent hydrogen bonding. a) Snapshot of a hydrogen bond between Thr152 and Asp162 from the ff14sb^fix^ trajectory at pH 7. b) pH-dependent deprotonation fraction of Asp162. c) pH-dependent occupancy of the hydrogen bond (h-bond) formation between the carboxylate of Asp162 and the hydroxyl group of Thr152. A h-bond is considered formed if the distance between the donor and acceptor heavy atoms is below 3.5 Å and the donor*−*H*···*acceptor angle is greater than 150^*°*^. d) Number of water molecules within 3.4 Å of a carboxylate oxygen of Asp162 at different pH. The data based on the ff19sb and ff14sbfix force fields are shown in red and black, respectively, and do not include p*K*_a_ corrections.

### p*K*_a_ downshift of His166 is due to solvent sequestration

To understand the significant p*K*_a_ downshift of His166, we examined its solvent exposure and hydrogen bond (h-bond) environment in the titration simulations (Figure 5a,b). Consistent with the analysis of the c22 titration simulations,^34^ the p*K*_a_ downshift of His166 can be mainly attributed to solvent displacement. Above pH 3.5, the first solvent shell of His166 includes only 3 water molecules, as His166 becomes protonated with decreasing pH, the number of water molecules in the first solvation shell of His166 increases from about 3 at pH 4 to about 5 at pH 1 (Figure 5c,d). Consistent with the c22 titration simulations,^34^ the h-bond formation of His166 is minimal. At pH 1–2, His166 (at N*δ*, Supplemental Figure S6) donates a h-bond to the backbone carbonyl oxygen of Asp162, with an occupancy of *∼*11% (Figure 5e).

**Figure 5:**
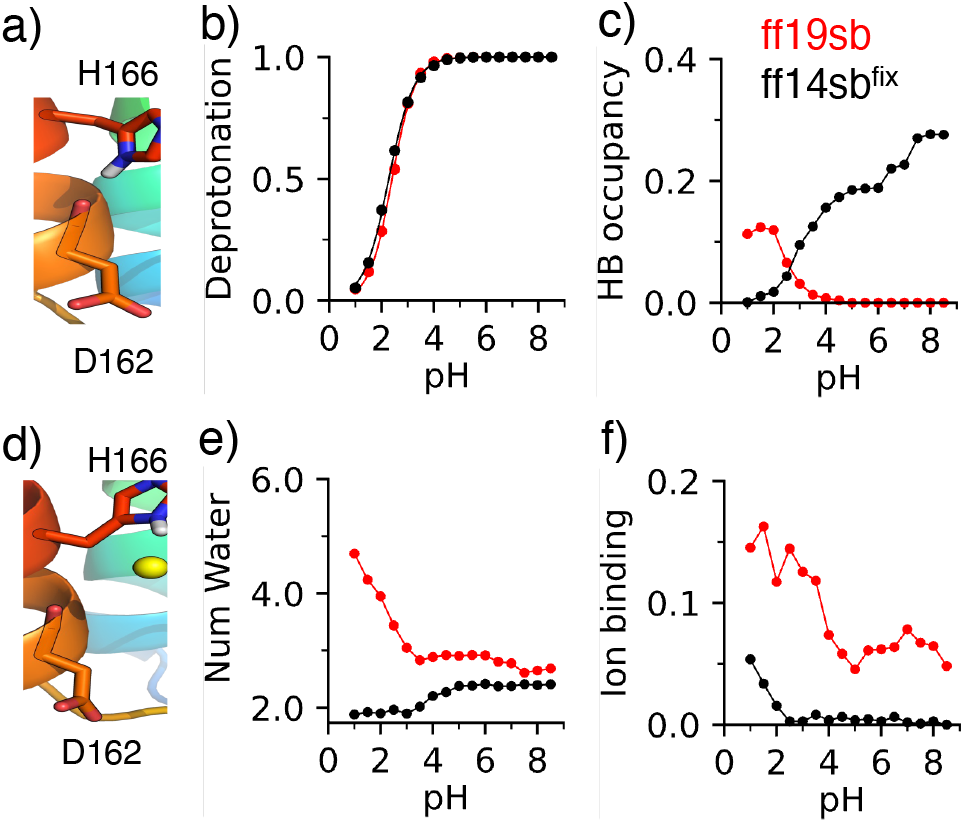
Titration of His166 in BBL and the pH-dependent hydrogen bonding, solvent exposure, and ion binding. a) Hydrogen bond formation between the imidazole nitrogen of His166 and the backbone carbonyl oxygen of Asp162 in a snapshot taken from the ff19sb trajectory at pH 2.5. b) The deprotonation fraction of His166 at different pH. c) Occupancy of the hydrogen bond (h-bond, shown in a) between His166 and Asp162. A h-bond is defined by a maximum distance of 3.5 Å between N and O and a minimum angle N*−*H*···*O of 150^*°*^. d) The imidazole nitrogen on His166 is bound to Cl^−^ (yellow sphere), shown in a snapshot taken from the ff19sb trajectory at pH 2.5. e) The number of water within 3.4 Å of an imidazole ni-trogen on His166 at different pH. f) The occupancy of Cl^−^ binding of His166 at different pH. A binding event is defined as a Cl^−^ ion within 4.2 Å of an imidazole nitrogen on His166. The data based on ff19sb and ff14sbfix are shown in red and black, respectively, and do not include p*K*_a_ corrections.

### NBFIX improves the p*K*_a_ calculations

To reduce the binding interactions between ions and proteins and between oppositely charged ions, atom-pair specific corrections to the Lennard-Jones parameters (commonly known as NBFIX) have been introduced to both CHARMM^47,48^ and Amber^49–51^ force fields. The ff19sb titration simulations showed specific binding of Cl^−^ with arginine and histidine and Na^+^ with glutamic acid (see below), and such interactions were not present in the c22 simulations^34^ with the NBFIX corrections.^48^

Since the NBFIX corrections for ff19sb have not been developed, we repeated the simulations of BBL using the NBFIX corrected ff14sb force field (ff14sb^fix^)^49–51^ and compared to the p*K*_a_’s calculated in the previous work^34^ based on the ff14sb force field, which gave a rmse of 1.78 or 1.38 without His166 (Supplemental Table S1). These errors are larger than those based on the ff14sb^fix^ simulations, which gave a rmse of 1.58 or 1.14 without His166 (Supplementary Table S1). It is worth noting that even with the NBFIX corrections, the p*K*_a_ errors based on ff14sb are still larger than the ff19sb simulations without the NBFIX corrections (rmse of 1.26 or 0.68 without His166). Below we discuss ion binding to the afore-mentioned residues, Glu141, Glu161, and Asp162, and His166 in the ff19sb simulations and the effects of using the NBFIX corrections.

### NBFIX abolishes ion binding and weakens salt bridges involving Glu141 and Glu161

In the case of Glu141, the ff19sb simulations showed a considerable amount of Cl^−^ binding to Arg137, which decreases from about 30% at pH below 4 to about 19% at pH 7.0 (Figure 2a and 2e). There is also occasional binding of a Na^+^ ion with Glu141, with a maximum occupancy of about 9% (Figure 2a and 2e). The introduction of NBFIX significantly weakened the interactions between Cl^−^ and Arg137 and between Na^+^ and Glu141, with the highest occupancy below 10% for either ion binding in the entire simulation pH range (Figure 2e). At the same time, the salt-bridge interaction between Glu141 and Arg137 is nearly abolished (Figure 2c), resulting in the increased solvent accessibility of Glu141 (Figure 2d). Comparing the p*K*_a_ of Glu141 based on ff14sb^fix^ and ff14sb, the 0.2-unit upshift can be attributed to the weakened salt bridge (Supplementary Table S1).

In the case of Glu161, the ff19sb simulations showed significant occupancies of chloride binding to Arg160 in the entire simulation pH range, which were abolished in the ff14sb^fix^ simulations (Figure 3f). At the same time, the salt-bridge interactions of Glu141 with Lys165 and Arg160 were also disrupted (Figure 3d), while the solvent accessibility was slightly increased (by half of a water, Figure 3e). We suggest that the abolishment of the salt-bridge interactions is the major reason for the 0.4-unit reduction in the p*K*_a_ down-shift for Glu161 in the ff14sb^fix^ as compared to the ff14sb simulations (Supplementary Table S1).

### The exaggerated p*K*_a_ downshift of Asp162 is reduced by the ff19sb relative to the ff14sb force field

As to Asp162, the ff19sb and ff14sb^fix^ simulations showed no ion binding or salt bridge formation. This explains why the calculated p*K*_a_’s based on the ff14sb^fix^ and ff14sb force fields are nearly identical. However, they are respectively 1.6 and 1.7 units lower than the value from the ff19sb simulations (Table 2 and Supplementary Table S1). The lower p*K*_a_ may be attributed to the stronger h-bond interaction with the hydroxyl group of Thr152, which stabilizes the deprotonated state of Asp162 (Figure 4c). Consequently, the pH profile of the h-bond occupancy is shifted to a lower pH range, coinciding with the pH range of the aspartic acid deprotonation (Figure 4b).

### NBFIX removes chloride binding with His166 and backbone h-bonding may be overstabilized by ff14sb

In the ff19sb simulations, a chloride ion occasionally binds His166 (mostly at the imidazole N*δ*), with an occupancy increasing from about 5% above pH 4 to over 15% below pH 2 (Supplementary Figure S5). The pH dependence of ion binding is inversely correlated with the histidine deprotonation, which is consistent with the electrostatic attraction between the positively charged histidine and Cl^−^ at low pH. The ff14sb^fix^ simulations abolished ion binding (Figure 5f), and increased the p*K*_a_ of His166 by 0.2 units relative to the ff14sb simulations (Supplementary Table S1). Interestingly, the p*K*_a_ based on ff19sb is 0.3 units higher than ff14sb^fix^ (Table 2). This may be attributed to the increased solvent accessibility, particularly below pH 4 where the number of water molecules is 5 in the ff19sb as compared to 2 in the ff14sb^fix^ simulations; this increased solvation stabilizes the charged state of histidine resulting in a higher p*K*_a_. Furthermore, the h-bonding interaction between His166 (exclusive to N*δ*) and the backbone carbonyl of Asp162 is negligible in the ff19sb simulations, whereas its occupancy increases with pH to nearly 30% at pH 8 in the ff14sb^fix^ simulations (Figure 5e). It is conceivable that the increased solvent exposure is due to the disruption of the sidechain-to-backbone h-bond.

## Concluding Discussion

In this work, we investigated the force field dependence of the all-atom PME CpHMD simulations using the p*K*_a_ calculations for a mini-protein which contains downshifted p*K*_a_’s of several carboxylic acids and one histidine. The CHARMM c22 and Amber ff19sb and ff14sb force fields were considered. Our data showed that the ff19sb and ff14sb force fields overestimate the p*K*_a_ downshifts of Glu141 and Glu161 involved in salt-bridge interactions and the p*K*_a_ downshift of the buried His166. These trends are consistent with the c22 force field but the overestimation by the ff19sb and ff14sb force fields is to a greater extent.

The p*K*_a_ data and pH-dependent conformational analysis suggest that the salt-bridge interactions involving Glu may be overly stabilized, which is a known deficiency of protein force fields.^29,32,51,65^ Application of the NBFIX corrections that significantly weaken ion binding and salt-bridge interactions in the ff14sb simulations^49–51^ demonstrated small upshifts of the calculated p*K*_a_’s; however, the reduction in the p*K*_a_ downshift remains insufficient. This is a topic that deserves further investigation in the future. Compared to the all-atom simulations, the p*K*_a_’s of Glu141 and Glu145 from the GBNeck2 simulations are much closer to experiment, although the p*K*_a_ downshifts remain slightly overestimated. This improvement may be in part explained by the larger Lys^+^–Glu^−^ and Arg^+^–Glu^−^ salt-bridge distances due to the difference in the geometry between GB and TIP3P.^60^

The overestimation of the p*K*_a_ downshift of His166 from the ff19sb and ff14sb simulations can be explained by the underestimated hydration free energy of the neutral histidine^31^ by the ff14sb force field and TIP3P or OPC3^66^ (similar to OPC^43^) water model, which suggests that the desolvation free energy of a buried histidine is too large. Interestingly, the p*K*_a_ of His166 is improved based on ff19sb relative to ff14sb, which may be attributed to the weakened intramolecular h-bonding and increased solvent accessibility (stronger solute-water interactions) in the OPC water.^43^ Compared to the Amber force fields, the hydration free energy of the neutral histidine based on the c22 force field and TIP3P model is closer to experiment, although that of the Hid tautomer is also underestimated.^67^ The difference in the hydration free energies between the ff14sb and c22 force fields explains why the p*K*_a_ of His166 is shifted lower based on the ff19sb or ff14sb relative to the c22 simulations. In stark contrast to the all-atom CpHMD simulations, the p*K*_a_ of His166 is somewhat overestimated in the GBNeck2 simulations, which may be due to the increased solvent exposure as a result of larger conformational fluctuation in the GB-Neck2 solvent.^68^

The largest improvement between the ff19sb and ff14sb calculated p*K*_a_’s is for Asp162. While ff19sb gives a p*K*_a_ in agreement with experiment, the ff14sb or ff14sb^fix^ overestimates the p*K*_a_ downshift by 1.5 or 1.6 units. Similar to His166, the h-bond involving Asp162 is weakened in the ff19sb simulations, which may be attributed to solute-water interactions in the OPC water^43^

One caveat of this work is worth mentioning. In the past, we developed titratable ions^11^ or water^69^ to rigorously account for the fluctuation in system net charge during titration; however, such a scheme was not applied due to slowed convergence and instead PME plasma (i.e., a uniform background charge) was employed for neutralization. To accommodate pH replica exchange, the number of ions was determined at pH 7.5 and not adjusted for low pH conditions, where Asp and Glu become protonated resulting in a net positive charge. Hub et al. demonstrated^70^ that invoking plasma stabilizes a test charge in a hexadecane slab due to the artificial background charge density in the low-dielectric region. In the BBL simulations, negatively charged plasma (at low pH) would stabilize positively charged His166 (the only buried residue), shifting its p*K*_a_ higher. However, this effect is not observed, likely because the local plasma density is low and the local dielectric constant is substantially higher than typically assumed (*ϵ* = 4) for the protein interior. Note, the same observation was made in our previous work.^34^ Nonetheless, we demonstrated previously^16,34^ that a box size dependent systematic shift in p*K*_a_ values is present in PME-CpHMD simulations and can be corrected by accounting for the plasma offset potential for water.^71^ In this work, no finite-size corrections^16,34^ were made due to the use of very large simulation boxes, which was shown^34^ to make corrections unnecessary.

Taken together, this study confirms that the accuracy of p*K*_a_ calculations using constant pH simulations is dependent on the underlying protein force field and water model. Given sufficient sampling, for example, in the case of the mini-protein BBL, deviations between the calculated and experimental p*K*_a_’s are largely reflective of the limitations of the force fields/water models. Salt-bridge overstabilization and underestimation of the histidine neutral state hydration are two common deficiencies of the ff19sb, ff14sb and c22 force fields (combined with their respective water models), although the magnitude of such errors appears to be larger for the ff19sb or ff14sb force fields. The BBL data also confirms that ff19sb with OPC is more accurate than ff14sb with TIP3P in terms of the balance between intramolecular and intermolecular protein-water interactions. Our work represents an initial effort to understand the force field limitations with the goal of improving the accuracy of all-atom PME CpHMD simulations.

## Data Availability

Parameter and simulation input files are freely available on github.com/JanaShenLab/ CpHMD_ff_comparison.

## Acknowledgement

Financial support is provided by the National Institutes of Health (1R35GM148261 and R01CA256557).

## Notes

### Competing Interest Statement

The authors have declared no competing interest.

### Summary of Updates

Additional discussion in Results and Conclusion.

