## Supplementary information for "Force Field Limitations of All-Atom Continuous Constant pH Molecular Dynamics"

### Contents

**List of Tables** **S-2**

**List of Figures** **S-2**

### List of Tables

S1 Comparison of the calculated p$K_\mathrm{a}$'s of BBL using the all-atom PME CpHMD titration based on different Amber force fields$^a$ . . . . . . . . . . . . . . . . . . S-3

S2 Model solvent exposure . . . . . . . . . . . . . . . . . . . . . . . . . . . . . S-3

### List of Figures

S1 Unprotonated fractions of BBL with ff19SB . . . . . . . . . . . . . . . . . . . . S-4

S2 Titration curves of BBL with the ff19SB force field . . . . . . . . . . . . . . . S-4

S3 Unprotonated fractions of BBL with the ff14SB$^{\text{fix}}$ . . . . . . . . . . . . . . . S-5

S4 Titration curves of BBL with the ff14SB$^{\text{fix}}$ force field . . . . . . . . . . . . . . S-5

S5 H-bonding occupancy of Asp162 . . . . . . . . . . . . . . . . . . . . . . . . . S-5

S6 Titration, H-bonding, and ion binding of the His166 tautomers . . . . . . . . S-6

Table S1: Comparison of the calculated p$K_a$'s of BBL using the all-atom PME CpHMD titration based on different Amber force fields$^a$

| Residue | Expt | ff19sb | ff14sb$^{fix}$ | ff14sb | ff14sb/GB |
|---|---|---|---|---|---|
| D129 | 3.9 | 3.5 | 3.6 | 2.6 | 2.8 |
| **E141** | 4.5 | 3.4 | 3.1 | 2.9 | 4.1 |
| H142 | 6.5 | 6.6 | 6.5 | 6.1 | 6.9 |
| D145 | 3.7 | 3.3 | 2.2 | 2.3 | 2.6 |
| **E161** | 3.7 | 2.4 | 2.4 | 2.0 | 3.3 |
| **D162** | 3.2 | 3.3 | 1.7 | 1.6 | 3.2 |
| E164 | 4.5 | 3.6 | 3.4 | 3.2 | 4.0 |
| **H166** | 5.4 | 2.4 | 2.1 | 1.9 | 6.0 |
| maxe | | -3.0 | -3.1 | -3.3 | -1.1 |
| rmse | | 1.26 | 1.58 | 1.78 | 0.66 |
| rmse (w/o H166) | | 0.73 | 1.14 | 1.38 | 0.67 |

$^a$Maximal error (maxe) and root-mean-square error (rmse) are calculated with respect to the experimental data. The experimental data of BBL is taken from Ref.[S1,S2] ff19sb refers to the ff19SB protein force field[S3] with the OPC water model[S4] and Amber default ion force field.[S5] ff14sb refers to the ff14SB protein force field[S6] with TIP3P water model[S7] and the Amber default ion force field.[S5] ff14sb simulations were conducted for 16 ns per pH replica, together with other published studies in our previous work.[S8] The model p$K_a$ corrections were made as described in Methods. Due to the much smaller box (10 Å cushion space between the protein heavy atoms and the closest water oxygen in the box edges), a -0.7-unit finite-size correction were made for all three types of residues.[S8] ff14sb$^{fix}$ refers to the ff14SB protein force field[S6] with TIP3P water model[S7] and the NBFIX corrected ion force fields.[S9–S11] c22 refers to the CHARMM c22 protein force field[S12] with the CHARMM style TIP3P water model[S7,S12] and the NBFIX corrected ion force fields.[S13–S15] Column ff14sb/GB contains the previous data[S16] obtained from the pH replica-exchange titration simulations based on the ff14SB force field and GBNeck2 implicit-solvent model.[S17] Residues discussed in the main text are highlighted in bold font.

Table S2: Number of water molecules within the first solvation shell of Asp, Glu, or His from the model penta-peptide simulations

| Residue | Num Wat |
|---|---|
| Asp | 6.99 ± 0.04 |
| Glu | 6.99 ± 0.06 |
| His | 4.94 ± 0.07 |

The number of waters within 3.4 Å of the carboxylate oxygen atoms on Glu, and Asp, or within the same distance from the imidazole nitrogen atoms in histidine. These results are averaged over the three ff19SB trajectories of the penta-peptide for the most exposed pH of each residue, i.e. pH 5.5 for Asp; pH 6.0 for Glu; pH 4.5 for His.

### Titration Convergence

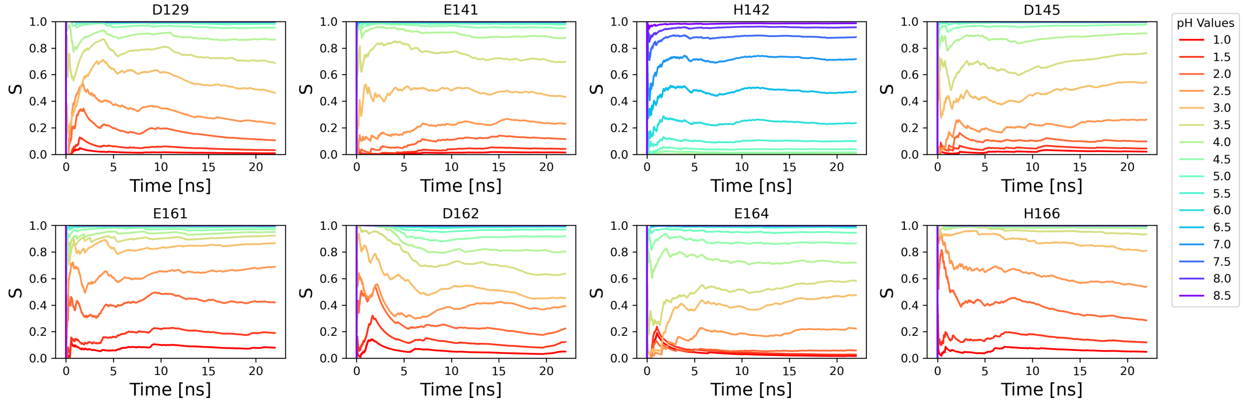


Figure S1: Time series of the unprotonated fraction of titratable residues in BBL from the simulations with the ff19SB force field.

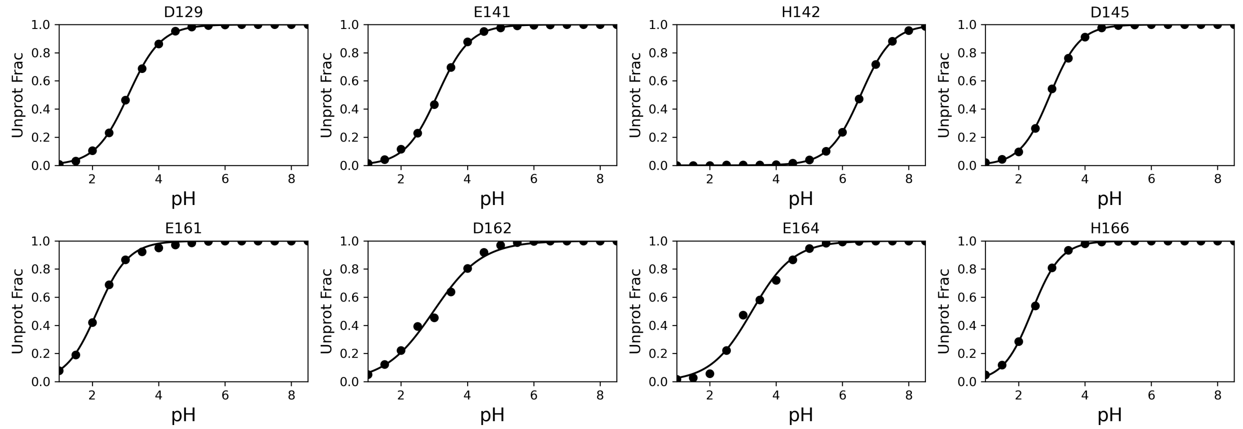


Figure S2: Deprotonation fractions vs. pH for the titratable residues from the simulations with the ff19SB force field.

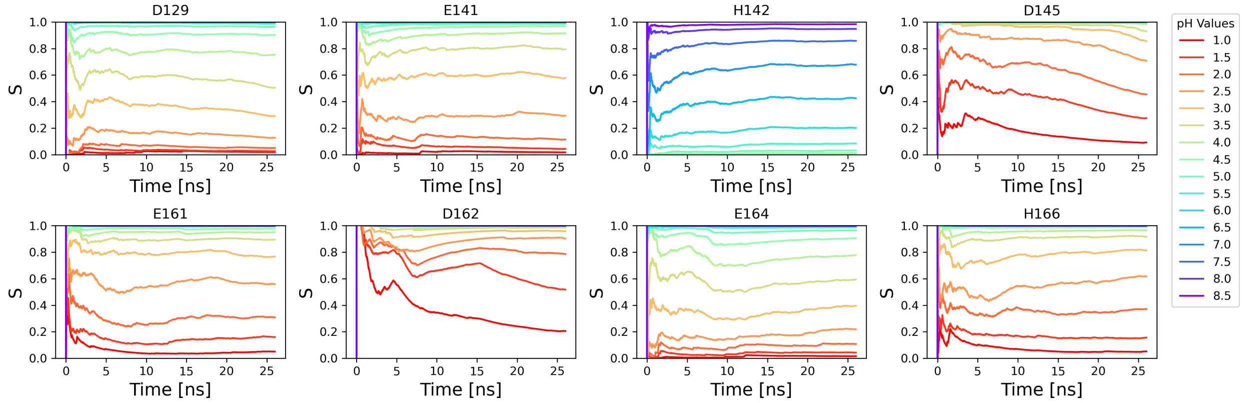


Figure S3: Time series of the unprotonated fraction of titratable residues in BBL from the simulations with the ff14SB$^{fix}$ force field.

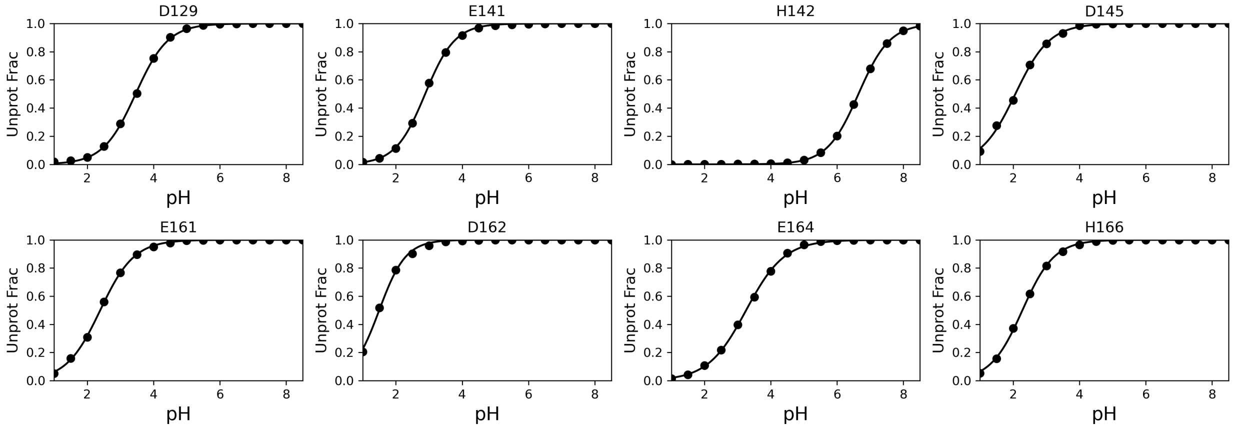


Figure S4: Deprotonation fractions vs. pH for the titratable residues from the simulations with the ff14SB$^{fix}$ force field.

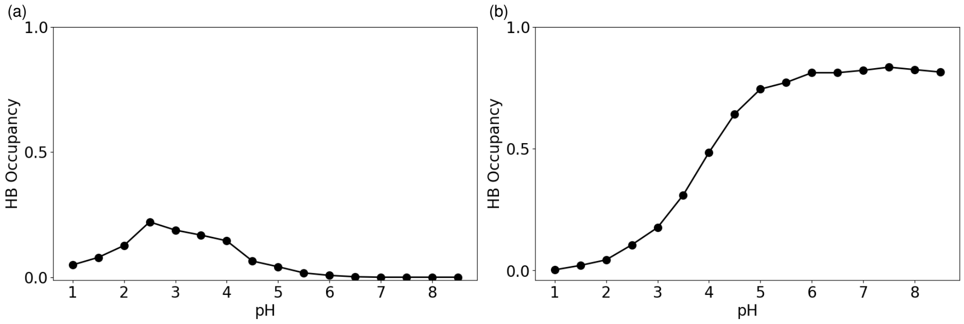


Figure S5: Hydrogen bonding between Thr152 and Asp162. pH-dependent occupancy of the hydrogen bond between T152 and protonated Asp162 (a) or deprotonated Asp162 (b).

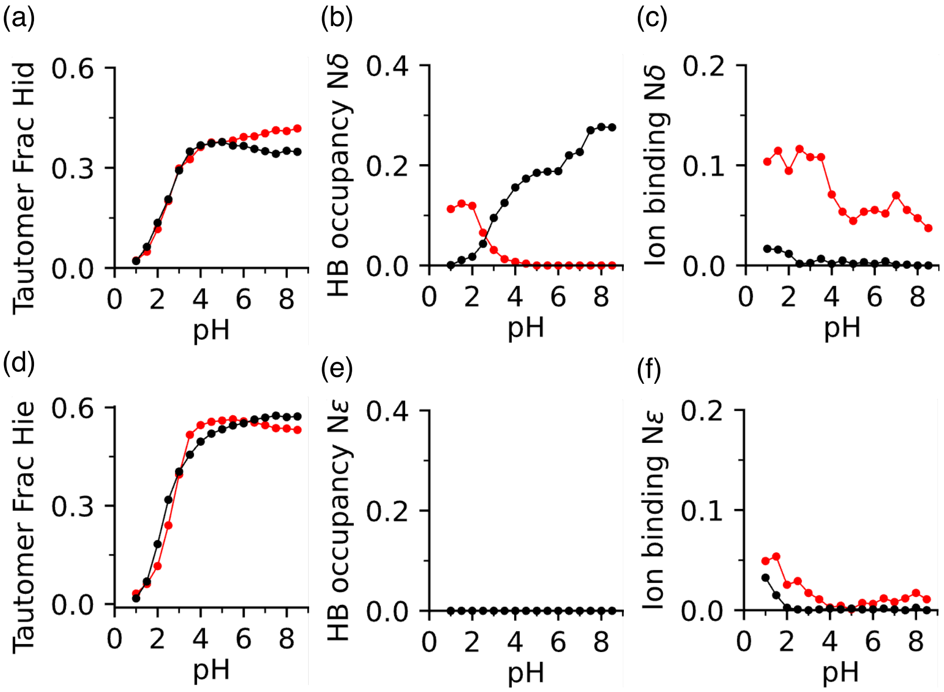


Figure S6: Hydrogen bonding and ion binding of His166 tautomers in the ff19SB and ff14sb$^{fix}$ simulations. a-c) pH-dependent fractions of the Hid tautomer (hydrogen on N$\delta$) and occupancies of the h-bond formation and ion binding at N$\delta$. d-f) pH-dependent fractions of the Hie tautomer (hydrogen on N$\epsilon$) and occupancies of the h-bond formation and ion binding at N$\epsilon$.

(S2) Arbely, E.; Rutherford, T. J.; Neuweiler, H.; Sharpe, T. D.; Ferguson, N.; Fersht, A. R. Carboxyl pKa Values and Acid Denaturation of BBL. *J. Mol. Biol.* **2010**, *403*, 313–327.

(S3) Tian, C.; Kasavajhala, K.; Belfon, K. A. A.; Raguette, L.; Huang, H.; Migues, A. N.; Bickel, J.; Wang, Y.; Pincay, J.; Wu, Q.; Simmerling, C. ff19SB: Amino-Acid-Specific Protein Backbone Parameters Trained against Quantum Mechanics Energy Surfaces in Solution. *J. Chem. Theory Comput.* **2020**, *16*, 528–552.

(S4) Izadi, S.; Anandakrishnan, R.; Onufriev, A. V. Building Water Models: A Different Approach. *J. Phys. Chem. Lett.* **2014**, *5*, 3863–3871.

(S5) Joung, I. S.; Cheatham, T. E. Determination of Alkali and Halide Monovalent Ion Parameters for Use in Explicitly Solvated Biomolecular Simulations. *J. Phys. Chem. B* **2008**, *112*, 9020–9041.

(S6) Maier, J. A.; Martinez, C.; Kasavajhala, K.; Wickstrom, L.; Hauser, K. E.; Simmerling, C. ff14SB: Improving the Accuracy of Protein Side Chain and Backbone Parameters from ff99SB. *J. Chem. Theory Comput.* **2015**, *11*, 3696–3713.

(S7) Jorgensen, W. L.; Chandrasekhar, J.; Madura, J. D. Comparison of Simple Potential Functions for Simulating Liquid Water. *J. Chem. Phys.* **1983**, *79*, 926.

(S8) Harris, J. A.; Liu, R.; Martins de Oliveira, V.; Vázquez-Montelongo, E. A.; Henderson, J. A.; Shen, J. GPU-Accelerated All-Atom Particle-Mesh Ewald Continuous Constant pH Molecular Dynamics in Amber. *J. Chem. Theory Comput.* **2022**, *18*, 7510–7527.

(S9) Yoo, J.; Aksimentiev, A. Improved Parametrization of $Li^+$, $Na^+$, $K^+$, and $Mg^{2+}$ Ions for All-Atom Molecular Dynamics Simulations of Nucleic Acid Systems. *J. Phys. Chem. Lett.* **2012**, *3*, 45–50.

(S10) Yoo, J.; Aksimentiev, A. Improved Parameterization of Amine–Carboxylate and Amine–Phosphate Interactions for Molecular Dynamics Simulations Using the CHARMM and AMBER Force Fields. *J. Chem. Theory Comput.* **2016**, *12*, 430–443.

(S11) Yoo, J.; Aksimentiev, A. New Tricks for Old Dogs: Improving the Accuracy of Biomolecular Force Fields by Pair-Specific Corrections to Non-Bonded Interactions. *Phys. Chem. Chem. Phys.* **2018**, *20*, 8432–8449.

(S12) MacKerell, A. D.; Bashford, D.; Bellott, M.; Dunbrack, R. L.; Evanseck, J. D.; Field, M. J.; Fischer, S.; Gao, J.; Guo, H.; Ha, S.; Joseph-McCarthy, D.; Kuchnir, L.; Kuczera, K.; Lau, F. T. K.; Mattos, C.; Michnick, S.; Ngo, T.; Nguyen, D. T.; Prodhom, B.; Reiher, W. E.; Roux, B.; Schlenkrich, M.; Smith, J. C.; Stote, R.; Straub, J.; Watanabe, M.; Wiórkiewicz-Kuczera, J.; Yin, D.; Karplus, M. All-Atom Empirical Potential for Molecular Modeling and Dynamics Studies of Proteins. *J. Phys. Chem. B* **1998**, *102*, 3586–3616.

(S13) Beglov, D.; Roux, B. Finite Representation of an Infinite Bulk System: Solvent Boundary Potential for Computer Simulations. *J. Chem. Phys.* **1994**, *100*, 9050–9063.

(S14) Lev, B.; Roux, B.; Noskov, S. Y. Relative Free Energies for Hydration of Monovalent Ions from QM and QM/MM Simulations. *J. Chem. Theory Comput.* **2013**, *9*, 4165–4175.

(S15) Venable, R. M.; Luo, Y.; Gawrisch, K.; Roux, B.; Pastor, R. W. Simulations of Anionic Lipid Membranes: Development of Interaction-Specific Ion Parameters and Validation Using NMR Data. *J. Phys. Chem. B* **2013**, *117*, 10183–10192.

(S16) Huang, Y.; Harris, R. C.; Shen, J. Generalized Born Based Continuous Constant pH Molecular Dynamics in Amber: Implementation, Benchmarking and Analysis. *J. Chem. Inf. Model.* **2018**, *58*, 1372–1383.

(S17) Nguyen, H.; Roe, D. R.; Simmerling, C. Improved Generalized Born Solvent Model Parameters for Protein Simulations. *J. Chem. Theory Comput.* **2013**, *9*, 2020–2034.
